# A novel gapmer nanobiosensor for probing long noncoding RNAs (lncRNA) expression dynamics during osteogenic and adipogenic differentiation

**DOI:** 10.1101/2023.07.01.547353

**Authors:** Samantha Fasciano, Shuai Luo, Shue Wang

## Abstract

Long non-coding RNAs (lncRNA) are non-protein coding RNA molecules that are longer than 200 nucleotides. lncRNA plays diverse roles in gene regulation, chromatin remodeling, and cellular processes, influencing various biological pathways. However, probing the complex dynamics of lncRNA in live cells is a challenging task. In this study, a double-stranded gapmer locked nucleic acid (ds-GapM-LNA) nanobiosensor is designed for visualizing the abundance and expression of lncRNA in live human bone-marrow-derived mesenchymal stem cells (hMSCs). The sensitivity, specificity, and stability were characterized. The results showed that this ds-GapM-LNA nanobiosensor has very good sensitivity, specificity, and stability, which allows for dissecting the regulatory roles of cellular processes during dynamic physiological events. By incorporating this nanobiosensor with living hMSCs imaging, we elucidated lncRNA MALAT1 expression dynamics during osteogenic and adipogenic differentiation. The data reveals that lncRNA MALAT1 expression is correlated with distinct sub-stages of osteogenic and adipogenic differentiation.

## Introduction

Osteoporosis is a bone disease that is characterized by loss of bone mass and structural deterioration of bone tissue, which can lead to a decrease in bone strength that can increase the risk of bone fractures. It is estimated that approximately 200 million people in the world are affected by osteoporosis, and more than 9 million fractures occur each year [1]. In the United States, it is estimated that more than 10 million people have osteoporosis, and about 54 million more people have low bone mass due to aging, which places them at increased risk [2]. It is reported that the cost of treating osteoporosis-related fractures is growing, and the predicted cost will rise to approximately $25.3 billion by 2025 [3]. One of the popular treatments for osteoporosis is to enhance osteogenesis or inhibit bone resorption through drug-based agents. For example, bisphosphonates are the predominant drug to treat osteoporosis by promoting osteoclasts apoptosis and inhibiting bone resorption rate [1, 4]. However, the major disadvantage of drug-based agents is their side effects and lack of capability to regain the lost bone density. Thus, there is an urgent need for alternative therapeutic treatments that are able to counteract bone mass loss.

Mesenchymal stem cells (MSCs) have great potential for tissue engineering, regenerative medicine, and cell-based therapy due to their capacity for self-renewal and multipotency. MSCs can be isolated from various sources, including bone marrow, adipose tissue, placenta, umbilical cord, or umbilical cord blood, respectively [5]. Under certain chemical or biophysical stimulation, MSCs can be differentiated into various lineages, including osteoblasts, adipocytes, neurons, and chondrocytes [6, 7]. MSCs also possess various physiological effects, such as maintenance of tissue homeostasis, regeneration, and immunomodulatory properties, making them valuable for cell-based therapeutic applications [8]. It is also reported that impaired osteogenic ability leads to less mature osteoblast formation and more adipocyte formation, which could eventually result in osteoporosis [4]. Although MSCs have great potential for cell-based therapies for treating different diseases, including osteoporosis, MSCs based-clinical trials are currently limited due to inconsistent differentiation capacity. However, the success of MSCs-based cell therapy is highly dependent on MSCs’ fate commitment. The uncontrolled differentiation can lead to an undesired phenotype, which limits their applications for tissue repair or regeneration. Thus, how to precisely control MSCs differentiation into the desired lineages is critical for MSCs-based therapies. Over the last few decades, unremitting efforts have been devoted to understanding biochemical signals that regulate MSCs’ commitment. Based on these efforts, a number of chemical stimuli (e.g., small bioactive molecules, growth factors, and genetic regulators) have been identified in regulating MSCs lineage commitment, including bone morphogenetic protein (BMP), Wnt, and Notch1-Dll4 signaling [9–11]. Moreover, several studies provide evidence that mechanical cues, including shear, stiffness, and topography, electrical stimulation, and acoustic tweezing cytometry (ATC) [12–14], both direct and indirect, play important roles in regulating stem cell fate. Moreover, it has been shown that ECM and topography enhance hMSCs osteogenic differentiation by cellular tension and mechanotransduction of YAP activity [15–18]. Recent advances have shown that long non-coding RNA (lncRNA), a class of non-coding RNAs with more than 200 bp in length, plays an important role in stem cell maintenance and specific lineage commitment [19, 20]. For example, it is reported that lncRNA-LULC activates smooth muscle differentiation of adipose-derived MSCs by upregulation of BMP9 expression; lncRNA MALAT1 [21], MSC-AS1 [22], lncRNA-OG, H19 [23–25], HHAS1 [26], NEAT1 [27] promote osteogenic differentiation either through miRNA-related regulation or chromatin remodeling [28]; lncRNA HOTAIR, MEG3 [29–31], and ANCR [32] were reported to inhibit osteogenic differentiation through miRNA-related regulation [33]. The complexity of the lncRNA signaling network requires novel dynamic gene analysis techniques to elucidate their specific roles in osteogenic differentiation. Current approaches for analyzing gene expression, single-molecule RNA FISH, RNA-seq, and single-cell transcriptomics [34–38], are often limited because these techniques require physical isolation or cell lysis. Therefore, it is crucial to investigate lncRNA expression dynamics during MSCs differentiation to elucidate the unrecognized characteristics and regulatory mechanisms that govern MSCs’ fate commitment.

To address the unmet needs, we developed a double-stranded gapmer locked nucleic acid (ds-GapM-LNA) nanobiosensor to detect and monitor lncRNA MALAT1 expression in live cells during osteogenic and adipogenic differentiation. We first characterized this nanobiosensor and demonstrated lncRNA expression dynamics during hMSCs differentiation. By incorporating this nanobiosensor with live cell imaging, we performed dynamic tracking of hMSCs and gene expression profiles of individual hMSC during osteogenic and adipogenic differentiation. We further investigated the role of lncRNA MALAT1 in regulating hMSCs during osteogenic and adipogenic differentiation.

## Materials and Methods

### Cell culture and reagents

hMSCs were acquired from Lonza and cultured in mesenchymal stem cell basal medium (MSCBM) with GA-1000, L-glutamine, and growth supplements. According to the manufacturer, hMSCs were originally isolated from normal (non-diabetic) adult human bone marrow withdrawn from bilateral punctures of the posterior iliac crests of normal volunteers. The hMSCs used in the paper are from three different donors. hMSCs were cultured at 37℃ in a humidified incubator with 5% CO_2_ with medium change every three days. Cells were passaged using 0.25% Trypsin-EDTA (ThermoFisher) once they reached 80-90% confluency. hMSCs from passages 3-6 were used in the experiments.

To induce osteogenic or adipogenic differentiation, hMSCs were seeded in 12-well plates with a density of 2 x 10^4^ cells/mL with a volume of 500 μL. When the cells reached 80% confluency, the basal medium was replaced with osteogenic or adipogenic induction medium. The osteogenic and adipogenic induction mediums were changed every three days. Images were taken daily for up to 15 days of induction, respectively.

### ds-GapM-LNA probe design and preparation

A ds-GapM-LNA probe consists of a gapmer LNA donor and quencher complex with a length of 30- and 15-base pair of nucleotide sequence with LNA-DNA-LNA monomers, respectively, **Tab. S1**. The donor sequence was labeled with a fluorophore at the 5’ end for fluorescence detection. The quencher sequence was labeled with an Iowa Black Dark Quencher at the 3’ end to quench the green fluorescence of the donor. To design the gapmer LNA donor sequence, the full sequence of lncRNA MALAT1 was first acquired from GeneBank. The minimum free energy structure of the lncRNA was computed using the RNA Fold web server. The target sequence can thus be selected and optimized by checking loop specificity. The LNA donor sequence is complementary to the loop region of the target lncRNA structure. The binding affinity and specificity were optimized using the mFold server and NCBI Basic Local Alignment Search Tool (BLAST) database. All the gapmer LNA sequences and DNA sequences were synthesized by Integrated DNA Technologies Inc. (IDT).

To prepare the ds-GapM-LNA complex, the gapmer LNA donor and quencher were initially prepared in 1x Tris-ethylenediaminetetra acetic acid (EDTA) buffer (pH 8.0) at a concentration of 100 nM. The gapmer LNA donor and quencher were mixed and incubated at 95 ℃ for 5 minutes in a pre-heated heat block and cooled down to room temperature over the course of two hours. To optimize the quenching efficiency, the donor probe and quencher probe were prepared in a number of different ratios to obtain a high signal-to-noise ratio. All the experiments were prepared in 384-well plates with triplets. The quenching efficiency was evaluated by measuring fluorescence intensity using a microplate reader (BioTek, Synergy 2). The prepared ds-GapM-LNA complex can be stored in a refrigerator for up to 7 days. For intracellular uptake, hMSCs were transfected with ds-GapM-LNA complex using lipofectamine 2000 (ThermoFisher) at a concentration of 100 nM for 24 hours. In brief, the complex was prepared in opti-MEM and mixed with lipofectamine 2000 for 15 minutes. After incubation, the mixed solution was added to each well.

### MALAT1 siRNA

To silence lncRNA MALAT1 expression, hMSCs were seeded in 12-well plates and transfected with MALAT1 antisense probe (Qiagen) at a concentration of 50 nM (final concentration) using Lipofectamine RNAiMAX (Cat. #13-778-150, Invitrogen) transfection reagent following manufacturer’s instructions. A negative control antisense probe was used as a negative control. After 24 hours of transfection, the transfection medium was replaced with a fresh basal culture medium. To investigate the effects of silencing MALAT1 on osteogenic and adipogenic differentiation, osteogenic and adipogenic induction was initiated after 24 hours of MALAT1 knockdown.

### Cell proliferation Assay

The effects of MALAT1 siRNA on hMSCs proliferation were characterized with a cell proliferation assay (Cell Counting Kit-8, CCK-8 assay, Sigma Aldrich). hMSCs were seeded in a flat-bottom 96-well tissue culture plate at a concentration of 2000 cells per well with a volume of 100 µL. After 24 hours of incubation for cell attachment and stabilization, hMSCs were treated with control siRNA and MALAT1 siRNA, following the manufacturer’s instructions. After 24 hours of silencing, cells were cultured in basal, osteogenic, and adipogenic induction medium, respectively. After five days of induction, a CCK-8 reagent was added to each well (10 µL per well). A microplate reader (BioTek Synergy H1) was used to measure the absorbance values of the samples at 450 nm. The fluorescence intensity value was normalized for comparison.

### Alkaline phosphatase activity (ALP) staining

To quantitatively assess the osteogenic differentiation, alkaline phosphatase (ALP) staining was performed using ALP live stain (ThermoFisher). To perform AP live staining, hMSCs were treated with ALP live staining at a concentration of 10x stock solution and incubated for 30 minutes, as per the manufacturer’s instructions. After staining, cells were washed twice using a basal culture medium, and images were captured after staining.

### Imaging and statistical analysis

All the images were captured using a ZOE Fluorescent Cell Imager with an integrated digital camera (BIO-RAD). To ensure consistency, all images were taken with the same settings, including exposure time and gain. Data collection and imaging analysis were carried out using NIH ImageJ software. To quantify ALP enzyme activity, the mean fluorescence intensity of each cell was measured, and the background noise was subtracted. All cells were quantified within the same field of view, and at least five images were quantified for each condition. The experiments were repeated at least three times, and over 100 cells were quantified for each group. Results were analyzed using an independent, two-tailed Student t-test in Microsoft Excel, with p < 0.05 being considered statistically significant.

## Results

### Intracellular lncRNA detection in live hMSCs

A double-stranded gapmer LNA/DNA (ds-GapM-LNA) nanobiosensor was developed by incorporating a gapmer LNA/DNA strand into another LNA strand to form a double-stranded gapmer LNA/DNA probe, **Fig. 1A**. Unlike previously reported LNA probes [7, 39–41], we modified the detecting probe by changing the location of the LNA monomers and the length of the detecting sequence. Specifically, instead of using alternating LNA/DNA oligonucleotides, we employed a gapmer LNA design, where the LNA oligonucleotides are located at the two ends of the detecting strand of the nanobiosensor, **Tab. S1.** The gapmer LNA probe is a 30-base pair single-stranded LNA/DNA oligonucleotide sequence with a fluorophore (6-FAM (fluorescein)) labeled at the 5’. The gapmer LNA probe sequence is complementary to part of the target lncRNA MALAT1 sequence. The quencher probe is a 15-base pair single-strand LNA/DNA oligonucleotide sequence. In the presence of a quencher probe, the gapmer LNA probe will bind to the quencher probe to form a double stranded gapmer LNA complex. The fluorophore at the 5’ will be quenched due to the quencher’s quenching ability. Once the LNA complex is transfected into the cells, in the presence of lncRNA target sequence, the gapmer LNA probe is thermodynamically displaced from the quencher, and binds to specific target sequence, **Fig. 1B**. This LNA probe displacement is due to a larger difference in binding free energy between LNA probe to target lncRNA versus LNA probe to quencher sequence. Furthermore, this displacement permits the fluorophore to fluorescence, thus detecting lncRNA gene expression at the single cell level, **Fig. 1C**.

**Fig. 1.**
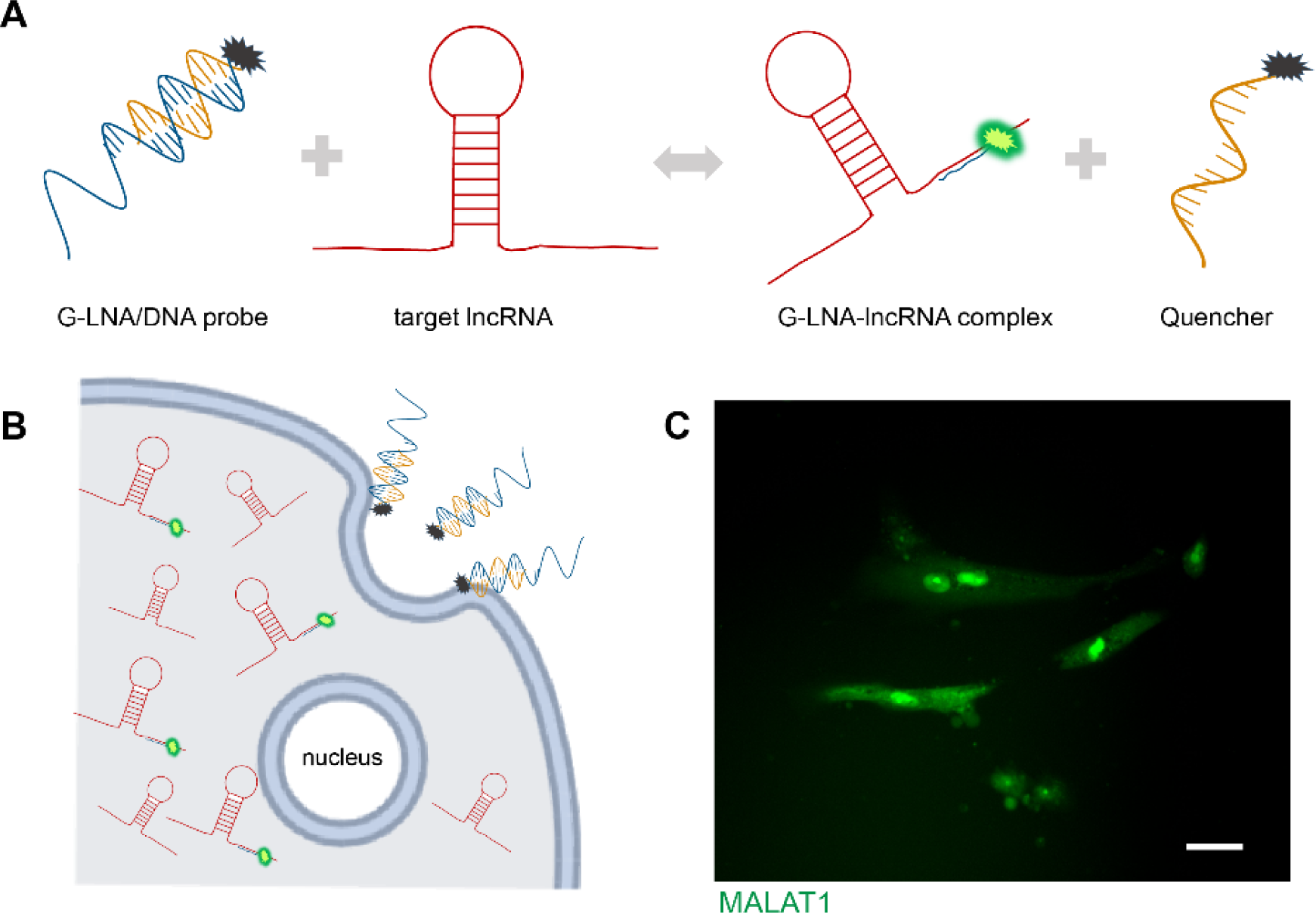
Working principle of ds-GapM-LNA nanobiosensor for detection of lncRNA in live cells. **(A)** Schematic illustration of this nanobiosensor. Briefly, this nanobiosensor is a complex of a gapmer LNA probe and a quencher probe. A fluorophore is labeled at the 5’ and is quenched due to close proximity. In the presence of target lncRNA sequence, the gapmer LNA probe is displaced from the complex and binds to the target sequence, allowing the fluorophore to fluorescene. **(B)** Schematic illustration of endocytic uptake of the nanobiosensor by cells for intracellular lncRNA detection. **(C)** Representative fluorescence image of lncRNA MALAT1 expression in single hMSC. Green: lncRNA MALAT1. Scale bar: 50 µm.

### Characterization of ds-GapM-LNA nanobiosensor

In order to optimize the ds-GapM-LNA nanobiosensor for monitoring lncRNA expression dynamics during hMSCs differentiation, we first characterized the optimal quencher-to-LNA ratio to minimize the background noise caused by free fluorophore during the reaction. The LNA probe concentration was set to 100 nM. The quencher-to-LNA probe ratio was then adjusted to make sure the quencher-to-LNA probe ratio ranges from 1 to 5. The fluorescence intensity was then measured at different quencher-to-LNA ratios, **Fig. 2A**. As expected, the fluorescence intensity decreases as the quencher-to-LNA probe ratio increases. To acquire the optimal ratio to minimize the background noise, the quenching efficiency at different quencher-to-LNA probe ratios was calculated. The quenching efficiency was determined by subtracting the fluorescence intensity of the free fluorophore from the fluorescence intensity of the LNA-quencher complex, divided by the intensity of the free fluorophore, multiplying the result by 100. Thus, the quenching efficiency was calculated as 63.7%, 82.7%, 94.8%, 97.5%, and 97.8%, as the quencher-to-LNA ratio was 1, 2, 3, 4, and 5, respectively. This result indicates that the quenching efficiency increases as the quencher-to-LNA probe ratio increases. The fluorescence intensity was quenched to a very low level, about 5% of the maximum intensity, at a quencher-to-LNA probe ratio of 3. Further increase in quencher concentration did not significantly increase the quenching efficiency. Thus, we set the ratio to 3 for the subsequent studies. This is different from the dsLNA/DNA probe, where the quenching efficiency was ∼97% when the quencher-to-donor ratio was 2 [7, 14, 39]. This result indicates the kinetic reaction between LNA donor and quencher depends on the length of the probe and the location of LNA monomers. We next characterized the detectable range of this ds-GapM-LNA nanobiosensor. We first prepared the LNA-quencher complex at a ratio of 3 to minimize the background noise. We next measured the fluorescence intensity by varying the DNA target oligonucleotide concentrations while the LNA probe concentration was set to 100 nM. As shown in **Fig. 2B**, the sigmoid-shaped titration curve shows an expansive dynamic range for quantifying lncRNA concentrations ranging from 1 to 1000 nM. Since the lncRNA concentration is typically low in mammalian cells (hundreds of copies) [42], this ds-GapM-LNA nanobiosensor is sufficient to detect target lncRNA at different concentrations. Furthermore, we evaluated the stability of this ds-GapM-LNA nanobiosensor by incubating the nanobiosensor with the target sequence for different durations, ranging from 3 to 15 days. As shown in **Fig. S1**, fluorescence intensity remained consistent regardless of the length of incubation time, suggesting that the stability of the nanobiosensor is not affected by the incubation duration.

**Fig. 2.**
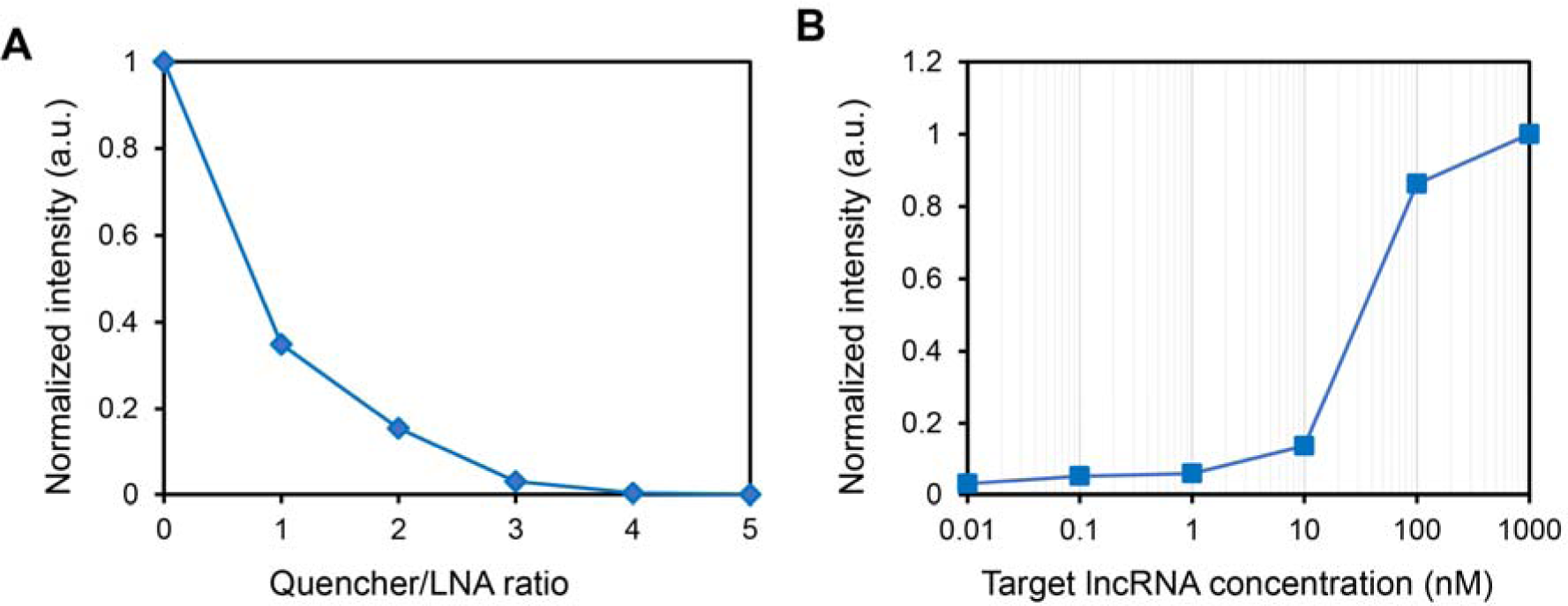
Characterization of ds-GapM-LNA nanobiosensor. **(A)** Optimization of the quencher-to-LNA probe ratio. Fluorescence intensity was measured by adjusting the quencher concentration using a microplate reader. The LNA probe concentration was set to 100 nM. **(B)** Calibration of the dynamic detection range of ds-GapM-LNA nanobiosensor by varying the lncRNA target concentration. The concentration of the LNA probe was set to 100 nM. The fluorescence intensity was normalized for comparison. All the experiments were performed in 384-well plates. Data are expressed as mean ±SEM. All the experiments were repeated at least three times independently with triplets.

The sensing performance of this ds-GapM-LNA nanobiosensor was evaluated and compared in hMSCs. lncRNA MALAT1 expression in individual hMSCs was detectable, **Fig. 3A**. A random GapM-LNA probe was used as a negative control. To evaluate the specificity of this ds-GapM-LNA nanobiosensor, MALAT1 expression was silenced using siRNA knockdown. A negative control siRNA was used for comparison. After siRNA treatments, hMSCs were transfected with random or MALAT1 probes to detect specific gene expression. For the random probe, there was no significant fluorescence signal with or without siRNA treatments, **Fig. 3B**. For the MALAT1 probe, siRNA treatment resulted in a reduction of MALAT1 expression when compared with negative control siRNAs, **Fig. 3C**. The knockdown efficiency was approximately 33.3%. It is noted that the fluorescence intensity, which indicates MALAT1 expression, was stable after 3, 5, and 7 days of incubation, **Fig. 3C**. Overall, this ds-GapM-LNA nanobiosensor displayed good performance in detecting intracellular lncRNA expression.

**Fig. 3.**
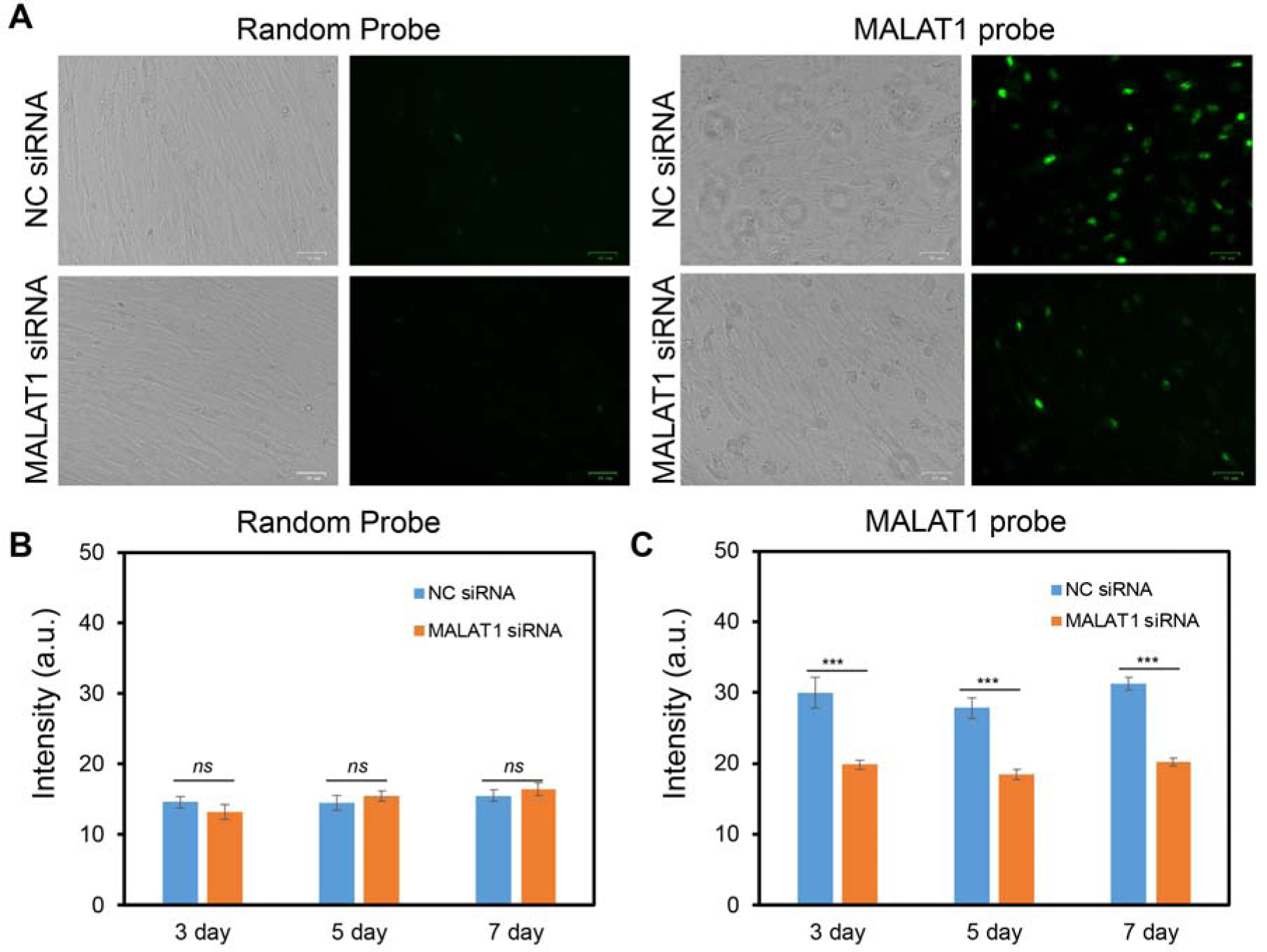
Detection of lncRNA MALAT1 using ds-GapM-LNA nanobiosensor. **(A)** Bright-field and fluorescence images of hMSCs with and without siRNA treatments. Scale bar: 50 µm. **(B)** Mean fluorescence intensity of random probe in hMSCs with and without MALAT1 siRNA knockdown. **(C)** Mean fluorescence intensity of MALAT1 expression with and without MALAT1 siRNA knockdown. Images are representative of at least three independent experiments. Data are expressed as mean ± s.e.m. (n = 3). p-Values were calculated using a two-sample t-test within groups. *, p < 0.05; **, p < 0.01; ***, p < 0.005.

### Dynamic gene expression analysis during osteogenic and adipogenic differentiation

Osteogenic and adipogenic differentiation are two dynamic cellular processes that consist of distinct sub-stages. Early differentiation involves distinct transcriptional responses as hMSCs differentiate into osteogenic or adipogenic lineages. Previous gene expression studies mainly focus on fixed cells, thus, the temporal information of gene expression dynamics is missing. It is crucial to understand how lncRNA expression relates to early lineage commitment. By employing this ds-GapM-LNA nanobiosensor, we monitored MALAT1 expression during hMSCs osteogenic and adipogenic differentiation. A random probe was used as a negative control. Both random and MALAT1 probes were transfected one day before the induction. MALAT1 expressions were monitored and measured daily for up to 15 days. The MALAT1 gene expression profile was determined by measuring the fluorescence intensity of individual cells using NIH ImageJ software. We confirmed the random probe has minimum fluorescence background in hMSCs during osteogenic and adipogenic differentiation. As shown in **Fig. 4B**, the fluorescence intensity of the random probe remained consistently low and did not exhibit any significant difference between these two groups on different days. Compared to random probes, MALAT1 expression exhibited different profiles during osteogenic and adipogenic differentiation, **Fig. 4C**, **Fig. S2-S4**. For osteogenic differentiation, MALAT1 expression showed a slow increase during the first week of differentiation and a greater increase during the second week of differentiation. In contrast, MALAT1 expression showed a slight increase in the first five days, and exhibited a rapid increase on day 7, indicating a distinct sub-stage transition during adipogenic differentiation. After seven days of differentiation, MALAT1 expression decreased gradually and eventually remained at a stable value on day 13 and day 15. These results indicate lncRNA MALAT1 expression follows different dynamic profiles during osteogenic and adipogenic differentiation. Especially, MALAT1 is highly expressed in the late stage (after seven days of induction) of osteogenic differentiation, but in the early stage (first seven days of induction) of adipogenic differentiation.

**Fig. 4.**
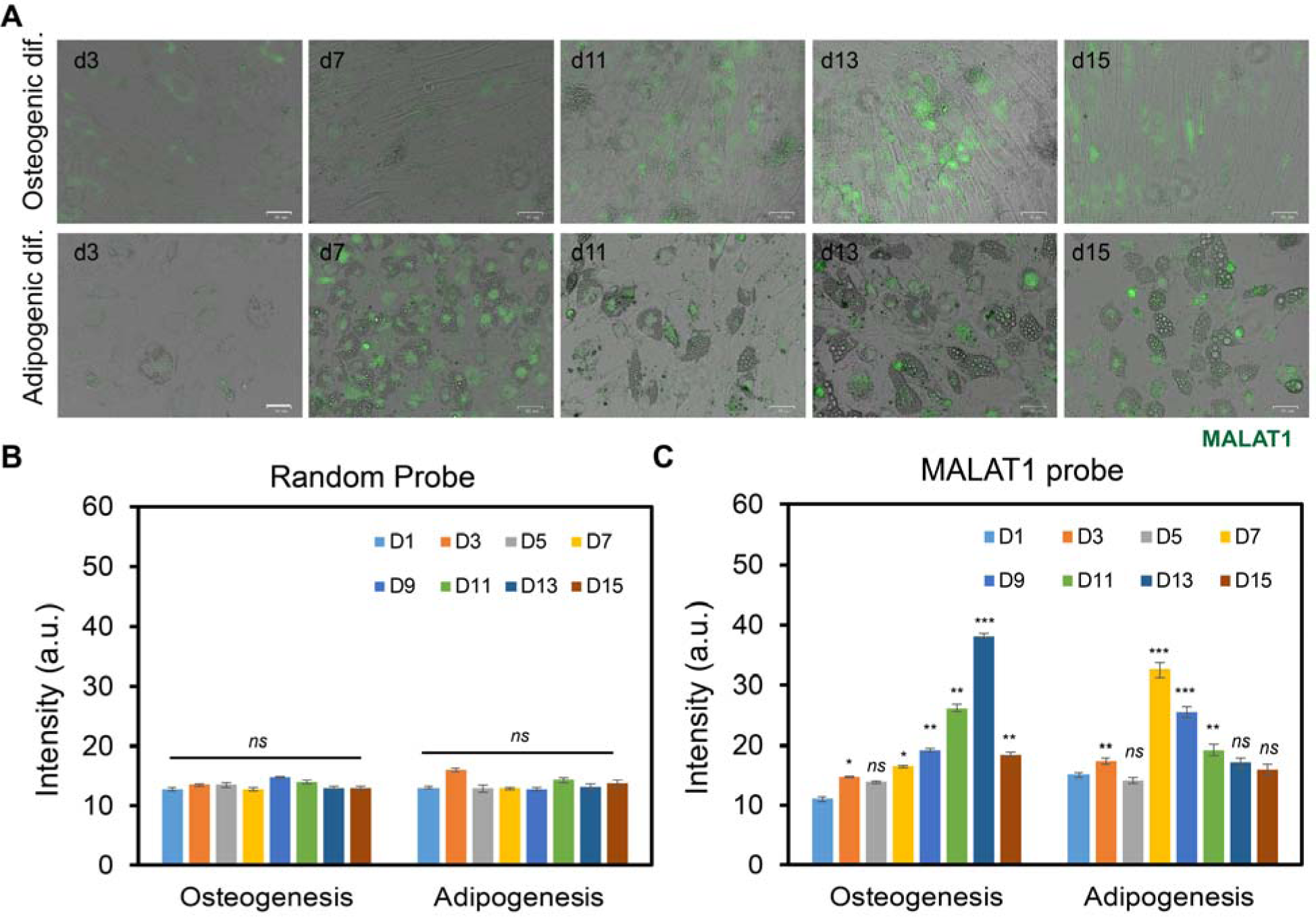
Dynamic MALAT1 expression analysis during osteogenic and adipogenic differentiation. **(A)** Representative merged images of hMSCs after 3, 7, 11, 13, and 15 days of osteogenic (upper panel) and adipogenic (bottom panel) differentiation. Green fluorescence indicates lncRNA MALAT1 expression. Scale bar: 50 µm. **(B)** Mean fluorescence intensity of random probe in hMSCs during osteogenic and adipogenic differentiation. **(C)** Mean fluorescent intensity of lncRNA MALAT1 in hMSCs at different time points during osteogenic and adipogenic differentiation. Images are representative of at least three independent experiments. Data represent over 100 cells in each group and are expressed as mean ± s.e.m. (n = 3). p-Values were calculated using a two-sample t-test with respect to day 1. *, p < 0.05; **, p < 0.01; ***, p < 0.005.

### MALAT1 expression indicates distinct sub-stages of osteogenic and adipogenic differentiation

Previous studies have indicated that there are distinct sub-stages of cell fate determination during both osteogenic and adipogenic differentiation. For osteogenic differentiation, there are three stages: early lineage progression (proliferation), early differentiation, and later stage differentiation (calcium mineralization). Early lineage progress involves differentiation initiation (0 – 3 hr), acquiring lineage (4-24 hr), and early lineage progression (24-48 hr). Early-stage differentiation occurs in the first five days, and later-stage differentiation occurs after five days of induction [7, 43]. In contrast, adipogenic differentiation comprises several distinct stages that can be characterized by specific cellular events: (1) commitment of hMSCs to adipocyte fate; (2) mitotic clonal expansion and proliferation, (3) lipid droplet accumulation, and (4) maturation and acquisition of adipocyte phenotype [44, 45]. To elucidate the correlation between MALAT1 expression and hMSCs osteogenic and adipogenic differentiation, we closely examined individual cells on various days to analyze the expression of MALAT1, **Fig. 5A-5B**. To identify the difference between the early stage (first seven days) and late stage of differentiation (after seven days), we took 1, 3, 5, 7, and 15 days as an example to compare MALAT1 expressions. For osteogenic differentiation, MALAT1 expression of these representative cells slightly increased after three days of induction and decreased at day 5. An increase reoccurred after seven days of induction and increased at day 15, **Fig. 5C**. This result indicates MALAT1 expression is involved in early osteogenic lineage progression and early-stage differentiation. It also indicates the heterogeneity of the hMSCs. For adipogenic differentiation, MALAT1 expression indicates different sub-stages of adipogenic differentiation. At the beginning, after adipogenic induction, cells go through growth arrest, with morphology change; this normally happens in the first 24 hrs. After growth arrest, cells went through mitotic clonal expansion (MCE) with gradually increased lipids. This process normally takes 2-4 days. After clonal expansions, cells go through early differentiation. At this stage, cells are committed to adipogenic differentiation. Terminal differentiation starts at nine days. At this stage, the nucleus is packed in the corner of a mature adipocyte, and the cell contains one large lipid vacuole, **Fig. 5B**. Interestingly, MALAT1 expression was increased during growth arrest, clonal expansion, and lipid accumulation. **Fig. 5D** demonstrates that the expression of MALAT1 decreases as lipid accumulation initiates. These findings suggest that MALAT1 expression could serve as a molecular indicator or signature of adipogenic differentiation.

**Figure 5.**
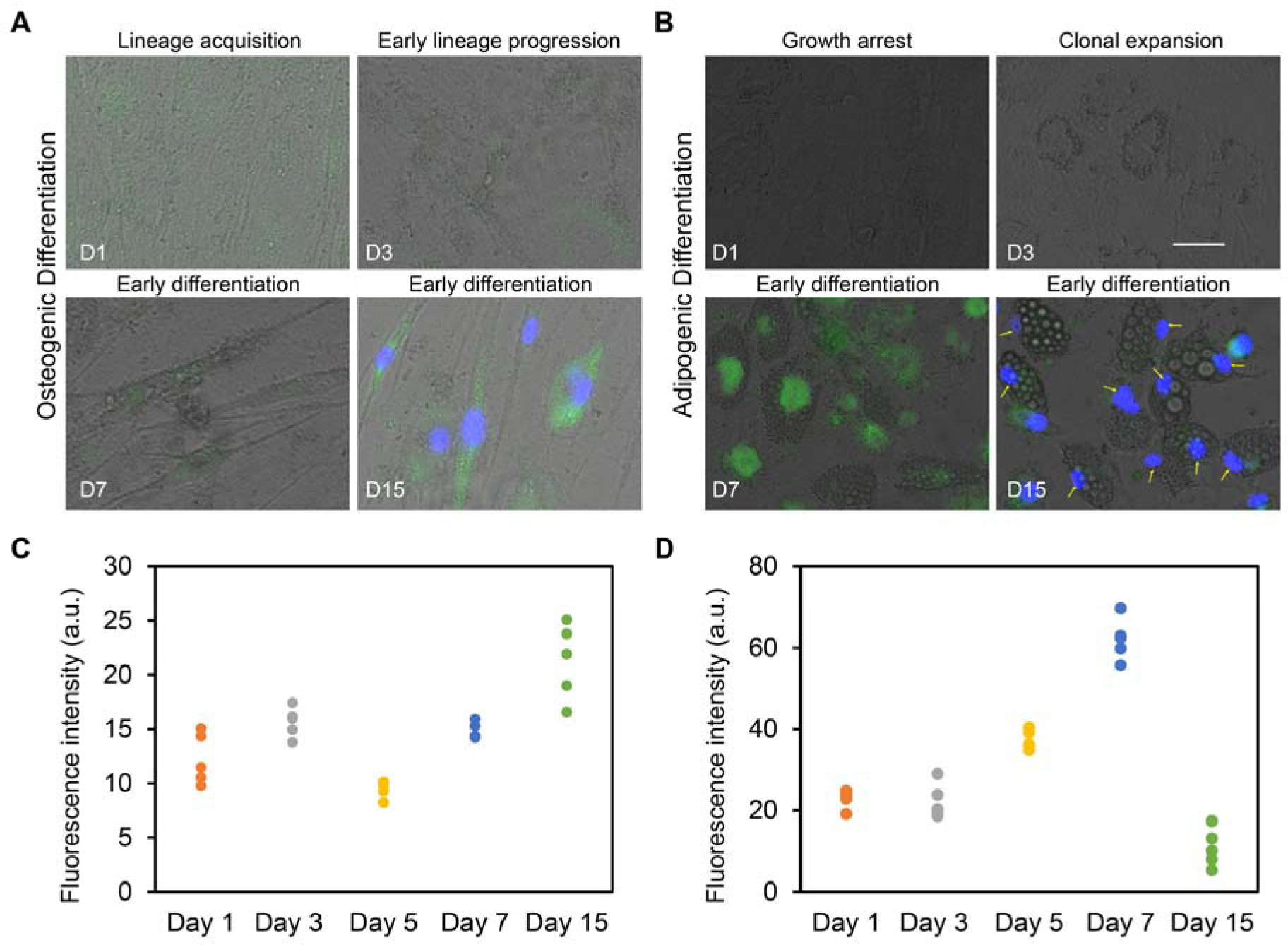
lncRNA MALAT1 expression distribution in individual cells. **(A)** and **(B)** are representative merged bright field and fluorescence field images at different days of osteogenic **(A)** and adipogenic **(B)** differentiation. Green fluorescence indicates MALAT1 expression. Yellow arrows indicate reduced MALAT1 expression after lipid accumulation during adipogenic differentiation. Scale bar: 50 µm. **(C)** and **(D)** are MALAT1 expression in individual hMSCs during osteogenic and adipogenic differentiation, respectively.

### MALAT1 regulates osteogenic and adipogenic differentiation

To investigate the role of MALAT1 in osteogenic and adipogenic differentiation, we disrupted the MALAT1 expression in hMSCs using siRNA knockdown. A negative control siRNA was used for comparison. The osteogenic differentiation after induction under different treatments was evaluated and compared, **Fig. 6A**. The cell proliferation of hMSCs after control siRNA and MALAT1 siRNA treatment was measured and compared, **Fig. S5**. Osteogenic differentiation was further quantified and compared by tracking alkaline phosphatase (ALP, an early biochemical marker for bone formation) enzyme activity using ALP live staining assay. **Fig. 6A** shows fluorescence images of hMSCs after 11 days of osteogenic differentiation. Interestingly, MALAT1 siRNA enhanced osteogenic differentiation with increased ALP enzyme activity. The ALP activities were further quantified and compared by measuring the mean fluorescence intensity at day five and day 11, **Fig. 6C**. Compared to control siRNA, the fluorescence intensity of ALP activity of hMSCs treated with MALAT1 siRNA exhibited a significant increase after five days and 11 days of induction. Furthermore, we conducted a further investigation to explore the role of MALAT1 in adipogenic differentiation. Interestingly, in contrast to osteogenic differentiation, MALAT1 siRNA inhibited adipogenic differentiation with deceased lipid droplets and nodules, **Fig. 6B**. We further evaluated the differentiation efficiency of hMSCs by quantifying the number of lipid droplets. The differentiation efficiency was determined by calculating the ratio of the number of lipid droplets per field to the total number of cells per field, multiplied by 100. **Fig. 6D** showed the comparison of the differentiation efficiency of hMSCs after 5 and 11 days. This result showed a significant decrease in adipogenic differentiation when silencing MALAT1. Taken together, these results suggest that lncRNA MALAT1 is involved in both osteogenic and adipogenic differentiation, exerting distinct effects in each process.

**Figure 6.**
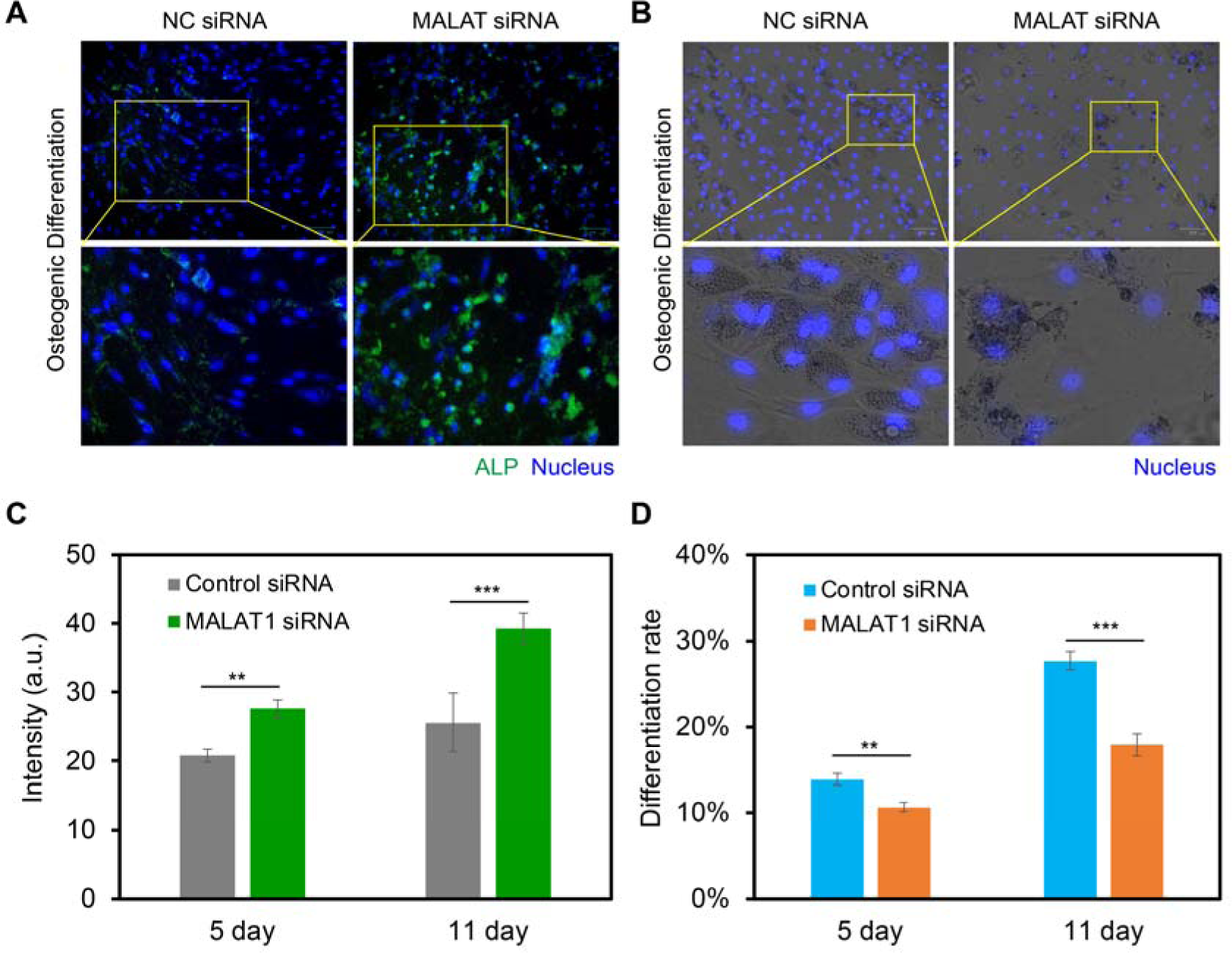
lncRNA MALAT1 in regulating osteogenic and adipogenic differentiation. **(A)** Representative fluorescence images of hMSCs during osteogenic differentiation with control siRNA and MALAT1 siRNA treatments. Green fluorescence indicates ALP enzyme activity. Blue indicates the nucleus. **(B)** Representative merged images of hMSCs during adipogenic differentiation with control siRNA and MALAT1 siRNA treatments. Blue indicates nucleus. Images were taken after 11 days of induction. **(C)** Mean fluorescence intensity of MALAT1 expression in hMSCs with control siRNA and MALAT1 siRNA knockdown. hMSCs were subjected to 5 days and 11 days of induction of osteogenic differentiation. **(D)** Comparison of hMSCs adipogenic differentiation efficiency under different treatments. Differentiation efficiency was determined by calculating the ratio of the number of lipid droplets per field to the total number of cells per field, multiplied by 100. Experiments were repeated independently at least three times. Data are expressed as mean± s.e.m. (n=4, ***, p<0.001, **, p<0.01)

## Discussion

In this study, we developed a novel ds-GapM-LNA nanobiosensor to detect lncRNA expression in hMSCs during osteogenic and adipogenic differentiation. This nanobiosensor, designed as a short sequence (30 nucleotides with LNA monomers at two ends), enables dynamic monitoring of lncRNA expression activities at the individual cell level. Unlike conventional lncRNA detection methods, this technique allows for dynamic gene expression analysis in live cells, eliminating the need for cell lysis or fixation. Compared to the previously reported dsLNA/DNA probe by our group [7], this new design demonstrated improved sensitivity, especially at low concentrations, **Fig. 2B**. This is crucial due to the low concentration of lncRNA in mammalian cells [42]. The specificity of this nanobiosensor was characterized and confirmed, in **Fig. 3**. Thus, this nanobiosensor exhibits high sensitivity, specificity, and stability, making it an ideal tool for tracking the dynamics of lncRNA during osteogenic and adipogenic differentiation. This ability enables us to establish a correlation between lncRNA MALAT1 expression and cell behaviors during osteogenic and adipogenic differentiation. The different MALAT1 expression correlates with cellular behaviors at different sub-stages in both osteogenic and adipogenic differentiation. Furthermore, this nanobiosensor demonstrates excellent resistance to nonspecific binding and maintains stability within the cells throughout the experimental duration. Thus, this ds-GapM-LNA nanobiosensor can be applied to study other dynamic cellular processes, including proliferation, migration, and differentiation.

lncRNA MALAT1, also known as NEAT2 (Nuclear-Enriched Abundant Transcript 2), is a highly conserved and abundant transcript in the nucleus with an extent of 8.7 kb. The role of MALAT1 is mostly seen in the nucleus, indicating a primary effect on gene transcription in collaboration with other regulators. It acts as a molecular scaffold, interacting with multiple proteins and influencing their localization and activity. MALAT1 has been shown to regulate gene expression at transcriptional and post-transcriptional levels, modulating chromatin organization, splicing, and mRNA stability. MALAT1 has been implicated in various physiological and pathological cellular processes, including cardiovascular disease [46], tumorigenesis, metastasis [47], osteoporosis [2], and stem cell differentiation [48–51]. Increasing evidence has shown that MALAT1 plays a crucial role in regulating osteogenic and adipogenic differentiation. Xiao et al. reported that MALAT1 was significantly increased during osteogenic differentiation of adipose-derived mesenchymal stem cells (ADSCs) [49]. This is consistent with our findings, which indicate that, MALAT1 expression gradually increased during hMSCs osteogenic differentiation, **Fig. 4**. Gao et al. found that MALAT1 could promote osteoblast differentiation of human bone marrow-derived mesenchymal stem cells (hBMSCs) from osteoporosis patients by targeting miR-143 [50]. Zheng et al. suggested that lncRNA MALAT1 expression was significantly reduced in osteoporosis rats compared with that of normal rats [51]. Furthermore, Yan et al. reported that inhibiting MALAT1 suppressed lipid accumulation and attenuated hepatic steatosis by reducing the stability of the nuclear SREBP-1c protein [52]. Another study showed that lncRNA MALAT1 knockdown inhibits the proliferation, migration, and tube formation of retinal endothelial cells [53]. Although numerous studies have demonstrated the involvement of lncRNA MALAT1 in osteogenic differentiation, the temporal MALAT1 expression in hMSCs during osteogenic and adipogenic differentiation has not been explored. Previous studies mainly focus on the effects of MALAT1 on cell differentiation instead of studying the dynamic gene expression analysis that is correlated to distinct stages of cell differentiation. Here, we studied the dynamic lncRNA MALAT1 expression during osteogenic and adipogenic differentiation of hMSCs. By exploiting a novel ds-GapM-LNA nanobiosensor, we studied the role of MALAT1 expression during early osteogenic and adipogenic differentiation. Our findings suggest that MALAT1 expression is correlated with distinct sub-stages of osteogenic and adipogenic differentiation. We also discovered MALAT1 plays different roles during osteogenic and adipogenic differentiation, respectively, **Fig. 4**. We further investigated the role of MALAT1 in regulating osteogenic and adipogenic differentiation by silencing MALAT1 expression in hMSCs. Our results indicate that knockdown MALAT1 results in increased osteogenic differentiation and decreased adipogenic differentiation. Further mechanistic studies are required to elucidate the molecular and cellular processes that are involved in MALAT1-regulated osteogenic and adipogenic differentiation. Specifically, the fundamental regulatory mechanisms of MALAT1 and its upstream and downstream signaling pathways should be further investigated using loss- and gain-of-function experiments.

## Conclusion

In this study, we developed a novel ds-GapM-LNA nanobiosensor to detect and monitor lncRNA MALAT1 expression during osteogenic and adipogenic differentiation of hMSCs. Unlike conventional lncRNA detecting techniques, such as RT-PCR, which requires a large number of cells, this ds-GapM-LNA nanobiosensor detects gene expression at the single-cell level. Another advantage of this nanobiosensor is its capability to study temporal gene expression dynamics during osteogenesis and adipogenesis without the requirement of cell lysis or fixation. Moreover, we demonstrated that this nanobiosensor exhibits high specificity, sensitivity, and stability. This non-invasive approach allows for real-time monitoring and analysis, providing valuable insights into the temporal changes of gene expression patterns throughout the osteogenic and adipogenic differentiation processes. Our study revealed that lncRNA MALAT1 expression is correlated with distinct sub-stages of osteogenic and adipogenic differentiation. Especially, MALAT1 expression was gradually increased during early osteogenic differentiation; while MALAT1 expression was rapidly increased once lipids accumulated during adipogenic differentiation. Knockdown of MALAT1 enhanced osteogenic differentiation while inhibiting adipogenic differentiation. In conclusion, with its high sensitivity, specificity, and stability, this nanobiosensor proves to be an ideal tool for tracking the dynamic changes of lncRNA during both osteogenic and adipogenic differentiation. Thus, this ds-GapM-LNA nanobiosensor can be utilized to investigate various dynamic cellular processes, such as proliferation, migration, differentiation, and development.

## Acknowledgment

S.W. would like to acknowledge the support from the NSF CMMI program (CAREER Award: 2143151). S. Fasciano is supported by the Provost Graduate Fellowship.

## Author contributions

S.W. conceived the initial idea of the study. S.L. and S.F performed the experiments. S.L., S.M, and S.W contributed to the experimental design and data analysis. S.L., S.M, and S.W. contributed to the writing. S.W. finalized the manuscript with feedback from all authors.

## Data Availability Statement

The original contributions presented in the study are included in the article/Supplementary Material; further inquiries can be directed to the corresponding authors.

## Notes

### Competing Interest Statement

The authors have declared no competing interest.

