## Supplemental Information for "A novel gapmer nanobiosensor for probing long noncoding RNAs (lncRNA) expression dynamics during osteogenic and adipogenic differentiation"

**Fig. S1.** Stability of ds-GapM-LNA nanobiosensor.

**Fig. S2.** Dynamic tracking of MALAT1 expression during osteogenic differentiation.

**Fig. S3.** Dynamic tracking of MALAT1 expression during adipogenic differentiation.

**Fig. S4.** Representative bright field and fluorescence images of hMSCs after 15 days under different treatments.

**Fig. S5.** Comparison of cell proliferation of hMSCs under control siRNA and MALAT1 siRNA treatments during osteogenic and adipogenic induction.

**Tab. S1.** ds-GapM-LNA probes and quencher sequences

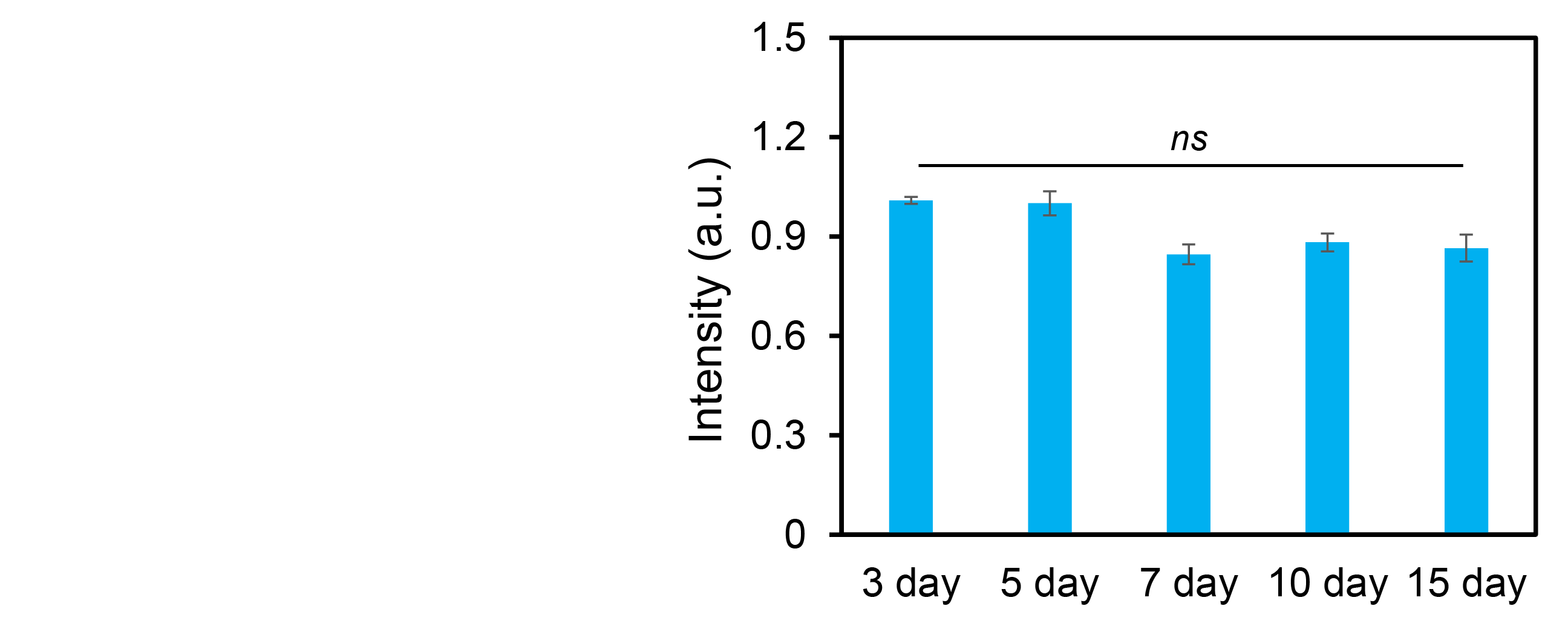

**Fig. S1**. **Stability of ds-GapM-LNA nanobiosensor**. Comparison of fluorescence intensity of ds-GapM-LNA nanobiosensor in the presence of target sequence. All the concentrations were set to 100 nM. Data are expressed as mean ±SEM. p-Values were calculated using a two-sample t-test within groups. *ns*, not significant.

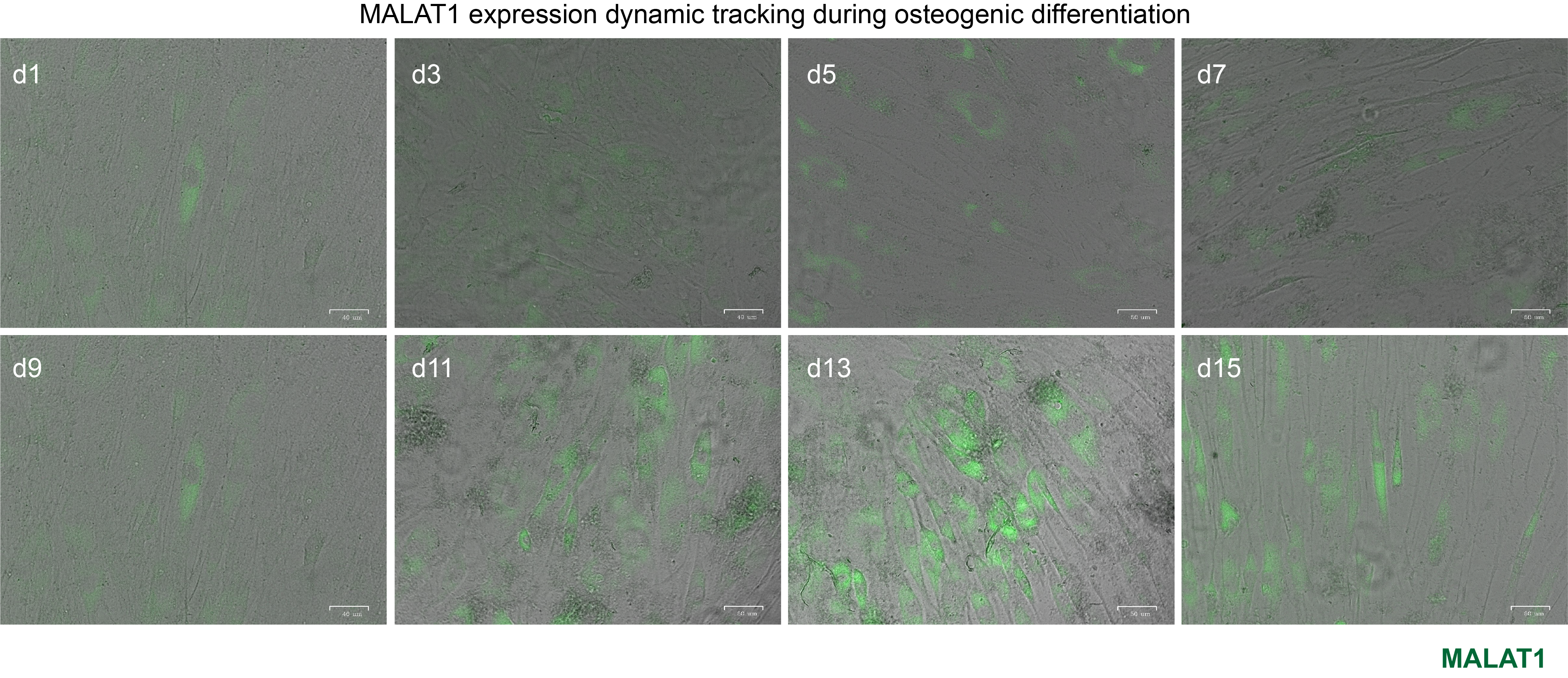

**Fig. S2. Dynamic tracking of MALAT1 expression during osteogenic differentiation.** Representative merged images of hMSCs during osteogenic differentiation. Images were taken every two days until 15 days of differentiation. Green fluorescence indicates MALAT1 expression. Scale bar: 100 µm.

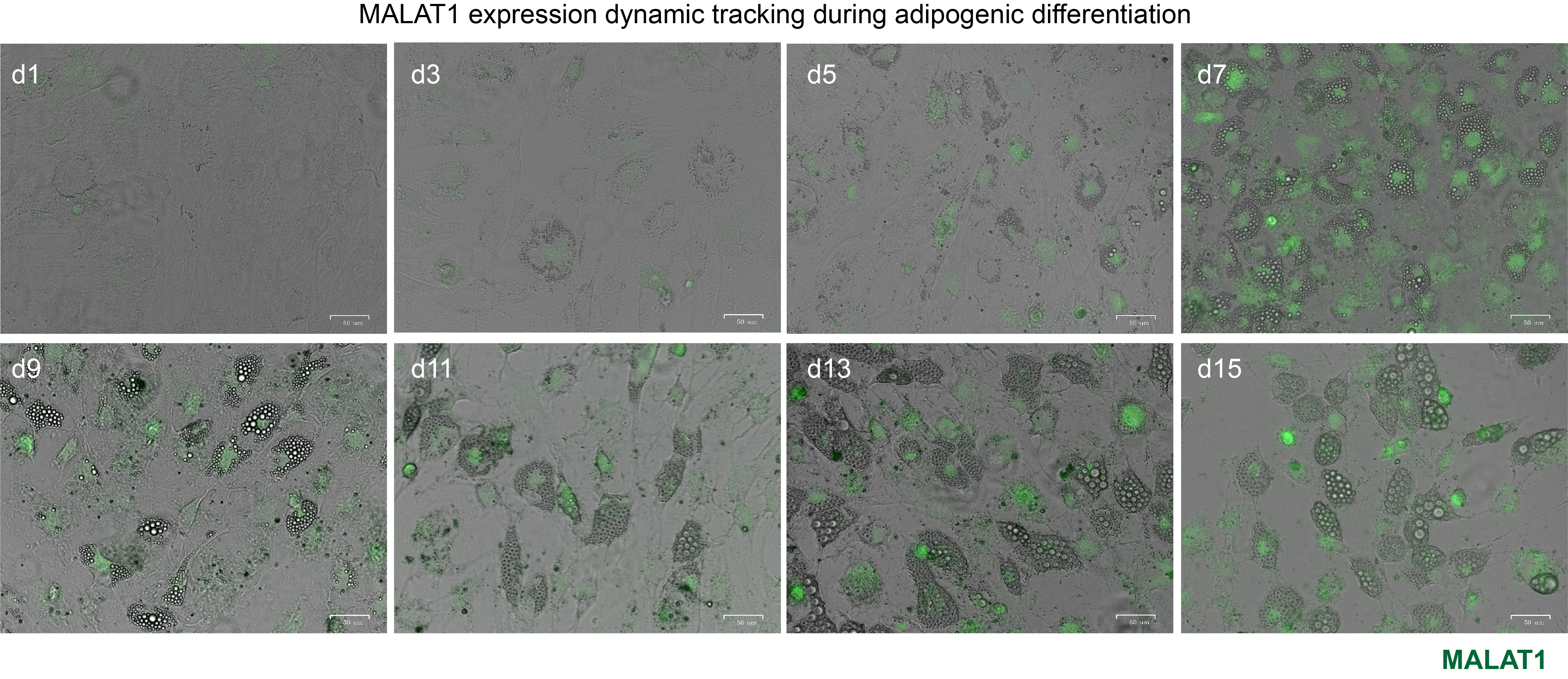

**Fig. S3. Dynamic tracking of MALAT1 expression during adipogenic differentiation.** Representative merged images of hMSCs during adipogenic differentiation. Images were taken every two days until 15 days of differentiation. Green fluorescence indicates MALAT1 expression. Scale bar: 100 µm.

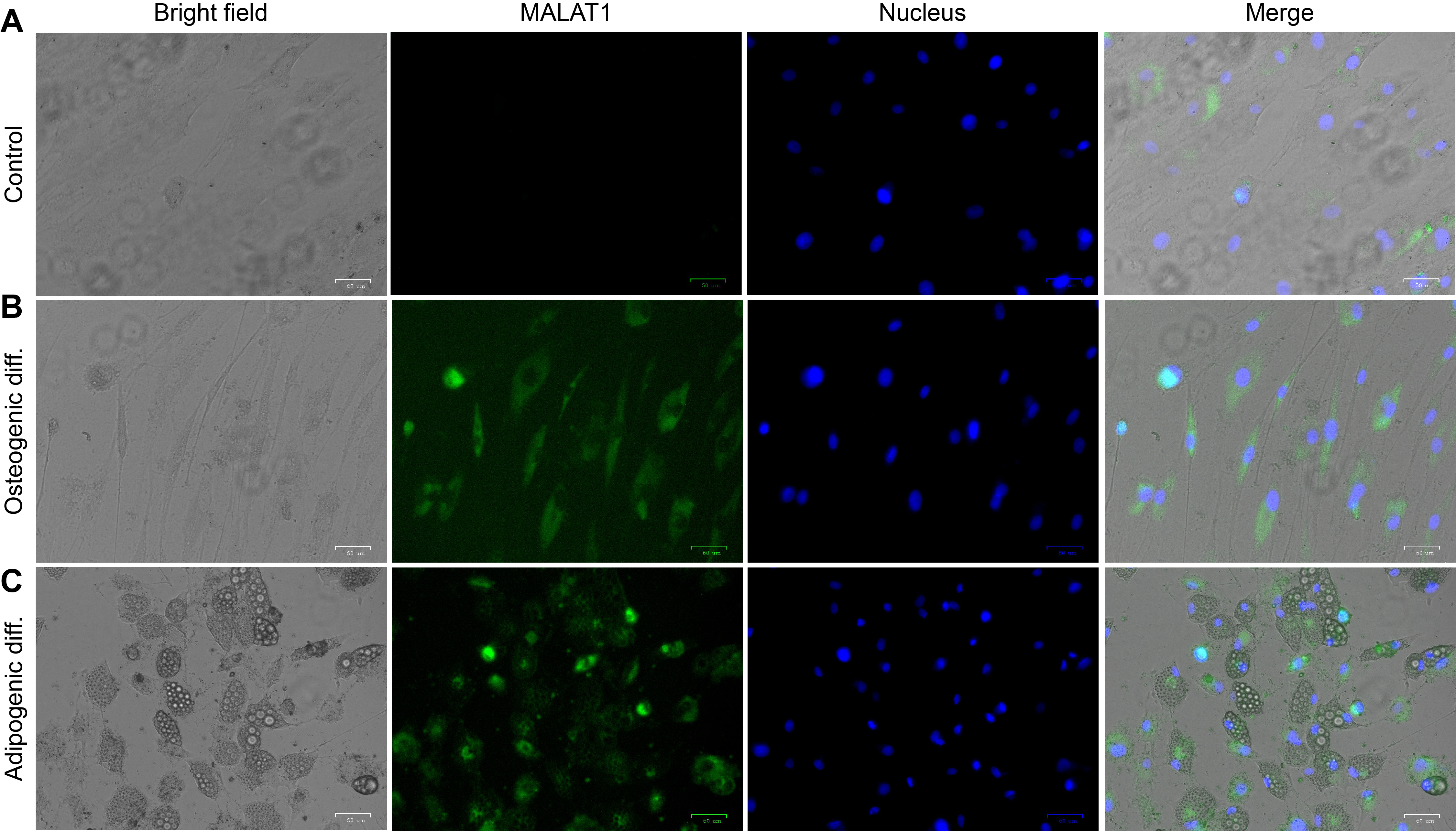

**Fig. S4.** **Representative bright field and fluorescence images of hMSCs after 15 days under different treatments.** **(A)** Control group; hMSCs were cultured in the basal medium without treatments. **(B)** Osteogenic induction group; hMSCs were cultured in osteogenic induction medium. **(C)** Adipogenic induction group, hMSCs were cultured in adipogenic induction medium. Green: MALAT1; Blue: Nucleus. Scale bar: 100 μm.

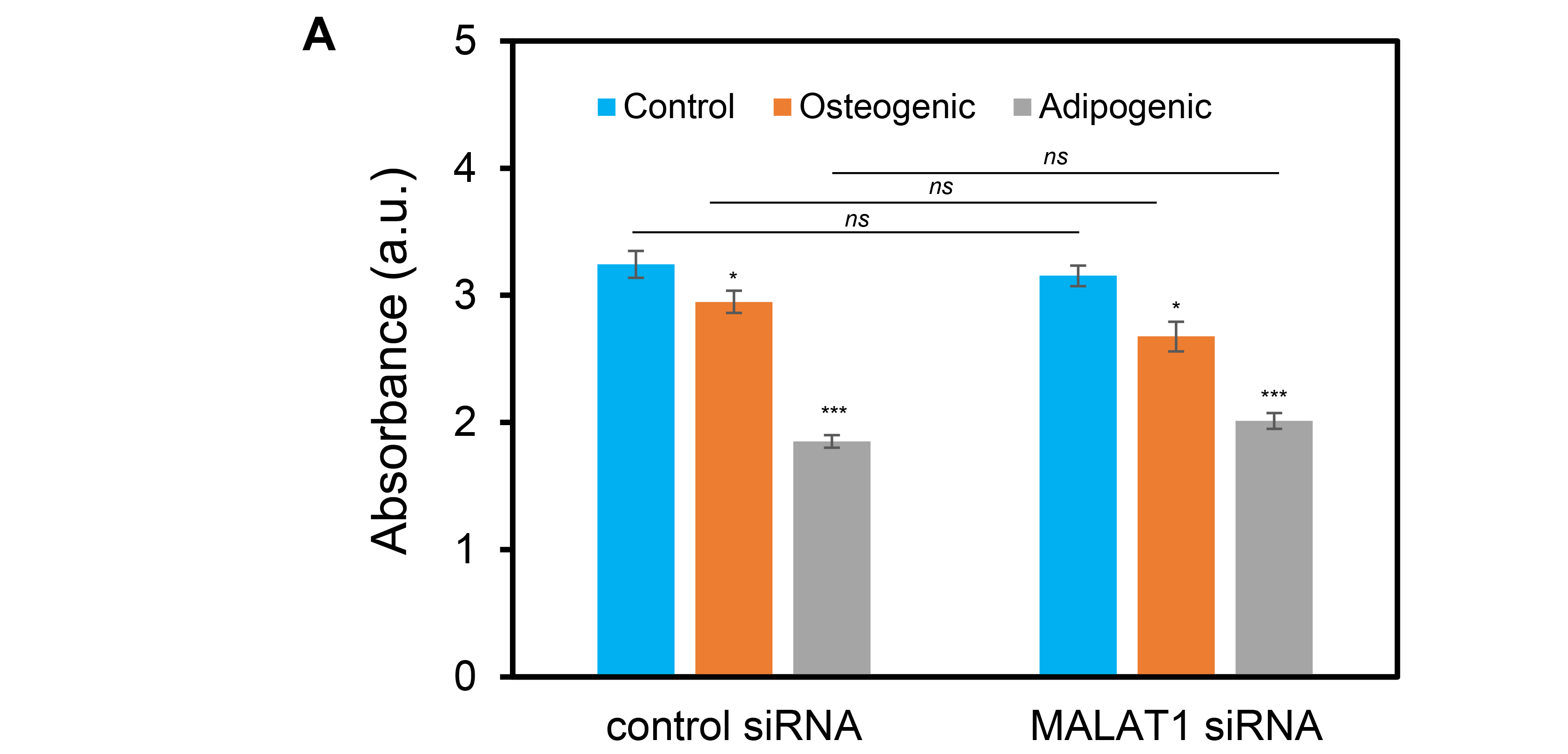

**Fig. S5**.  **Comparison of cell proliferation of hMSCs under control siRNA and MALAT1 siRNA treatments during osteogenic and adipogenic induction.**  For the control group, hMSCs were maintained in basal culture medium. hMSCs were seeded in 96-well plates with 2000 cells per well. After cells reached 80% confluency, hMSCS were treated with control siRNA and MALAT1 siRNAs. After 24 hours of silencing, hMSCs were induced to osteogenic or adipogenic differentiation. The cell proliferation was evaluated using a cell counting kit (CCK-8) assay. Absorbance was measured and compared at 450 nm. Experiments were repeated independently at least three times. Data are expressed as mean± s.e.m. (n=3, ***, p<0.001, **, p<0.01)

**Tab. S1.** ds-GapM-LNA probes and quencher sequences

| Name | | Sequence (5’-3’) | Fluorophore |
| --- | --- | --- | --- |
| Dll4 mRNA | Donor | +T+C+G+C+A TACGT GTGTC TGCTG AGTGT +T+C+C+T+G | /56-FAM |
|  | Quencher | +G+A+C+A+C ACGTA TGCGA | /3-Iowa BlackFQ |
|  | Target | CAGGA ACACT CAGCA GACAC ACGTA TGCGA |  |
| Random | Donor | +T+A+C+A+G TATCT CGAAG ACCAG TAG GG +C+A+C+C+T | /56-FAM |
|  | Quencher | +C+T+T+C+G AGATA CTGTA | /3-Iowa BlackFQ |
|  | Target | AGGTG CCCTA CTGGT CTTCG AGATA CTGTA |  |

* + represents LNA monomer
